# Microfluidic system-based time-course tracking of physical proximity between cells and its effect on gene expression for elucidating live single cancer-immune cell interactions

**DOI:** 10.1101/2021.11.13.468447

**Authors:** Bianca C. T. Flores, Smriti Chawla, Ning Ma, Chad Sanada, Praveen Kumar Kujur, Ludmilla T. D. Chinen, Kyle Hukari, Mark Lynch, Naveen Ramalingam, Debarka Sengupta, Stefanie S. Jeffrey

## Abstract

Cell-cell communication and physical interactions play a vital role in cancer initiation, homeostasis, progression, and immune response. Here, we report a system that combines live capture of different cell types, co-incubation, time-lapse imaging, and gene expression profiling of doublets using a microfluidic integrated fluidic circuit (IFC) that enables measurement of physical distances between cells and the associated transcriptional profiles due to cell-cell interactions. The temporal variations in natural killer (NK) - triple-negative breast cancer (TNBC) cell distances were tracked and compared with terminally profiled cellular transcriptomes. The results showed the time-bound activities of regulatory modules and alluded to the existence of transcriptional memory. Our experimental and bioinformatic approaches serve as a proof of concept for interrogating live cell interactions at doublet resolution, which can be applied across different cancers and cell types.

## Introduction

Cell-cell communication sustains the multicellular organism as an integral unit via direct physical interactions, surface receptor-ligand interaction, cell signaling from adjacent cells, nearby cells, or even distant organs^1,2^. The investigation of cell-cell interaction in the tumor microenvironment (TME) is one of the barriers to understanding cancer progression and identifying new therapeutic targets^3^. Despite the advances in high-throughput microscopy and single-cell based molecular analysis, tools to precisely quantify live cell-cell interactions are lacking or, more recently, characterized using combinations of spatial -omics techniques on fresh-frozen or formalin-fixed complex tissues and live cell analyses^4–8^. We developed a microfluidic workflow involving capture and co-incubation of live single stromal/cancer cells or doublets using the single-cell dosing mRNA-seq integrated fluidic circuit (IFC) system (Fluidigm^®^), which provides both spatial and transcriptional cell-cell interactions. To demonstrate the performance for the quantification of the cell-cell interaction, we applied our platform for natural killer (NK) - triple-negative breast cancer (TNBC) cancer-immune doublets (CIDs).

TNBC was chosen as a model due to its aggressive nature and lack of estrogen receptor, progesterone receptor, and human epidermal growth factor receptor 2 that limit the use of targeted therapies^9,10^. However, promising responses have been seen with immunotherapy^11^, which is emerging as an important component of cancer treatment^12,13^. Clinically promising discoveries exploit specific features of tumor-immune cell crosstalk involving immunosuppression and anti-tumoral response signaling^14–17^. Among immune cells, NK cells can efficiently kill multiple neighboring cells with oncogenic transformation of surface markers^18,19^. NK cell activation, combined with their capacity to enhance antibody responses, supports NK cells’ role as anticancer agents^20^. It is speculated that genetically engineered endogenous NK cells can exert tumor immunosurveillance and influence tumor growth^21,22^. However, heterogeneity is ubiquitous in human cancer making the selection of personalized treatment/therapy a challenge. It is expected that NK cell heterogeneity further contributes to NK-tumor crosstalk dynamics with differential modulation of their cytotoxic response, triggering tumor death when the balance between activation and inhibitory protein levels are considered^23–25^. It is essential, therefore, to characterize the molecular level cues emanating from single NK cells when they encounter tumor cells. Thus, a single-cell platform for NK-cancer cell interaction measurement is much needed to study NK-cancer immunotherapy.

To better understand NK-tumor cell interactions, we present a microfluidic workflow involving capture and co-incubation of single NK and cancer cells (CIDs) using the Polaris^TM^ Single-Cell Dosing mRNA Seq IFC^26^ (Fluidigm). The doublets captured in the microfluidic chambers were tracked for cell-cell distances, using time-lapse imaging. After 13 hours of incubation with the exchange of growth medium at a defined interval of time (5 hours), the cells were subjected to single-cell RNA sequencing (scRNA-seq). This offered a total of 290 transcriptomes, including single NK and cancer cells as control and NK-cancer cell doublets (CIDs). Unsupervised clustering analysis of the scRNA-seq expression profiles revealed heterogeneity in the TNBC cell line. We also observed a small number of NK killing events among the co-incubated CIDs, which allowed us to characterize the gene expression signature associated with NK-mediated lysis event of the companion cancer cells. We correlated the hourly computed cell-cell distances with terminally profiled gene expression vectors. We noted the existence of transcriptional memory, governed by precise regulatory modules active in a time-bound manner. In addition, we investigated the ligand-protein pairs interactions, which provided cues for the inflated activity of CD24/SIGLEC10 and ANXA1/EGFR in the tumor-NK doublets and supporting a previously described interaction between CD24/SIGLEC10 as a potent immunotherapy target for ovarian and triple negative breast cancer^27^.

## Results

To study the interactions between NK and TNBC cells, we captured single NK-92MI cells (interleukin-2 independent Natural Killer cell line), single MDA-MB-231 cells, and cancer-immune doublets (CIDs, one NK-92MI cell and one MDA-MB-231) and incubated the cells for 13 hours using the Fluidigm Polaris system^26,28–31^ (Fig. 1A, Fig S1). Time-lapse images of CIDs were captured every hour to measure the distance between the CIDs. Following incubation, single cells and doublets were processed for RNA-sequencing. The integrated fluidic circuit can perform on-chip multi step chemistry that includes cell capture, co-incubation, lysis, reverse transcription, and cDNA amplification^29^. Single-cell and CIDs were analyzed for expression of marker genes that distinguished the two cell types. We identified previously known markers for both NK-92MI cells and MDA-MB-231 cells. In the case of NK cells, we observed strong expression of NK cell marker genes *KLRD1*, *LAIR1*, *CCR6* and *TNFRSF9*. Single cancer cells showed TNBC marker genes *HMGA1*, *ANKRD11*, and *TACSTD2* (Fig. 1B) when analyzed using SCANPY^32^ toolkit for analyzing single-cell gene expression data. Further to validate our cell type annotations, we took advantage of the imaging capability of the Polaris system. Prior to cell selection on the microfluidic IFC, the NK and cancer cells were stained with CellTracker™ Deep Red Dye and CellTracker™ Orange CMRA Dye respectively. The z-score normalized intensities of NK and cancer cell channels were subjected to 2D visualization (plotted using R package scatterD3) revealing the grouping of cells as per their annotations based on cell staining (Fig S2). Unsupervised clustering with Seurat v3 and associated Uniform Manifold Approximation and Projection (UMAP) based 2D visualization^33^ of the transcriptomes revealed two separate clusters, which were primarily dominated by clonal heterogeneity of the cancer cell line (Fig. 1C). The single NK cells shared one of the clusters, cluster 1, with a cancer cell sub-group. Upon performing principal component analysis (PCA) exclusively on the transcriptomes from this cluster, we noted spatial segregation of the NK cells (Fig. 1C). To further delineate the cancer cell heterogeneity, we performed unsupervised clustering of single cancer cells separately, which also resulted in two distinct clusters (Fig S3A), featuring differentially expressed genes (Fig S3B).

**Fig 1.**
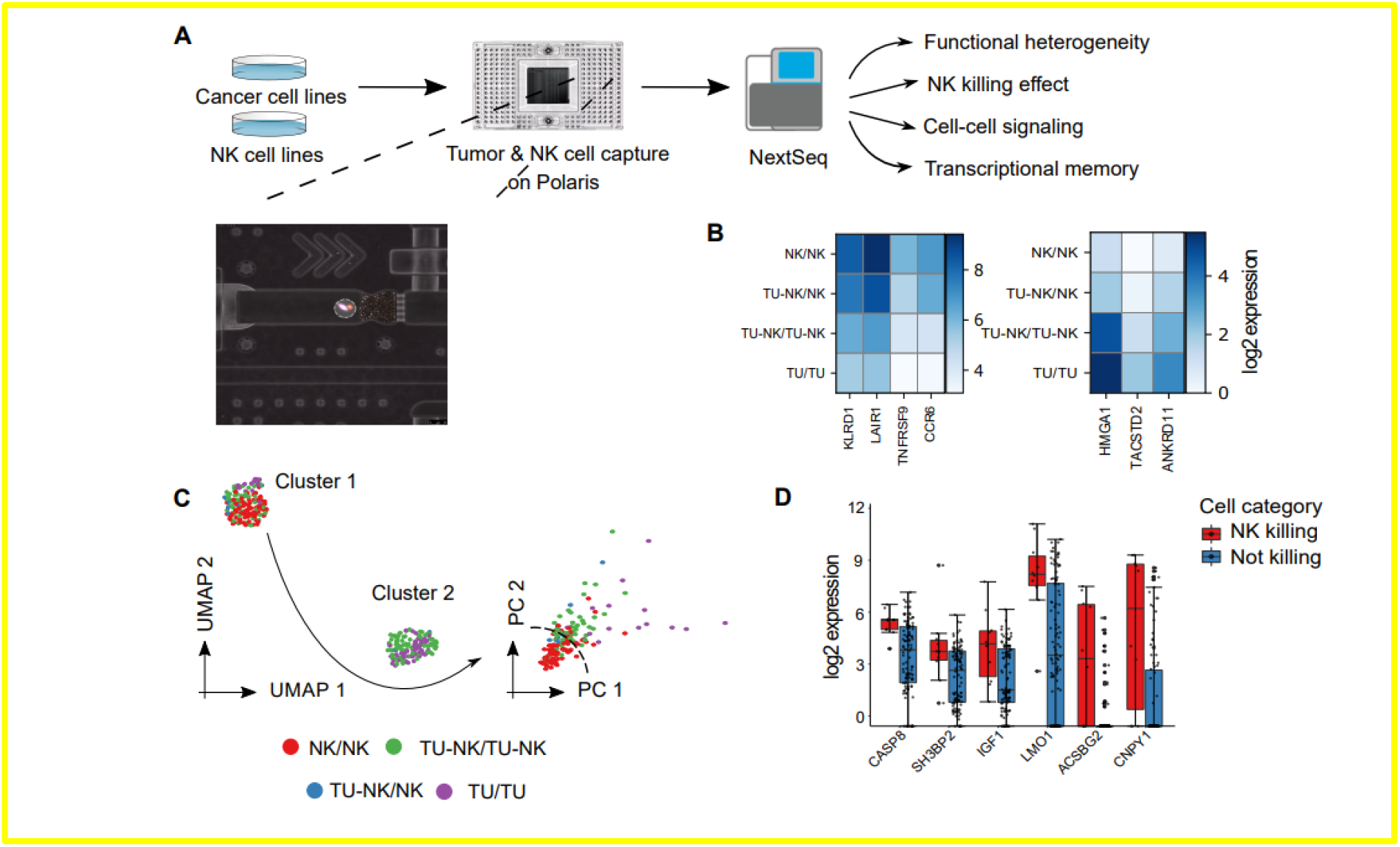
(A) Schematic workflow for cell interaction studies. The workflow involves cancer and NK cells propagate in culture and stained off-chip, which are then captured as live single cells and doublets, incubated/co-incubated an imaged over time. Inset shows a microscopic image of a chamber on the microfluidic Integrated Fluidic Circuit (IFC) containing an MDA-MB-231 (Blue) and NK cell (Red) doublet. The cells are then lysed, reverse transcribed, and the cDNA amplified *in-situ* within the chambers; down-stream library preparation, sample barcoding, and sequencing are performed off-chip using the Illumina NextSeq system. (B) Heatmap plotted using SCANPY showing average expression of marker genes for single NK and cancer cells, confirming their lineage identity. (C) UMAP-based visualization of the cells shows two separate clusters mainly due to tumor cell line heterogeneit (tumor cells in purple). A further dimension reduction of cluster 1 using PCA shows the separation of tumor (purple) and NK cells (red). Legend key indicates cell status at the beginning and the end of the time-course tracking (e.g., TU-NK/NK denotes interactions which initially consisted of both tumor and the NK cells in a chamber, an subsequently the NK cell remained in the chamber after co-incubation due to a cell killing event). (D) Boxplots showing selected differentially expressed genes distinguishing killing vs non-killing events.

### Distance Tracking of Cancer-Immune Cell Interactions Over Time Shows Transcriptional Memory

We tracked CIDs for dynamic changes in the distance between cancer and the companion NK cells under incubation in the same microfluidic chamber over 13 hours (Fig. 1A). At the end of the 13^th^ hour, transcriptome sequencing was performed on the CIDs (*n* = 102). This allowed us to determine the association between terminally measured gene expression with cancer/immune cell distances measured across different time points. Transcriptomes profiled at the end of the 13^th^ hour featured transcripts that correlate significantly with CID distance measurements across all the time points (Fig 2A, 2B). Surprisingly, we identified time-bound activities of at least three distinct gene modules (Fig 2B). At time point 6 (after five-hour incubation and a change in culture medium), we noted a new set of genes (module 2; M2) expressed. However, we did not observe a similar shift in gene expression at later culture medium change time points. We performed module wise transcription factor activity analysis using ShinyGO^34^ and RcisTarget^35^, which inferred the regulatory role of three potential transcription factors, namely BRCA1, YY1, and THAP1. Among these, BRCA1 is predicted by ShinyGO, whereas YY1 and THAP1 along with their candidate target genes by RcisTarget (Fig 2C).

**Fig 2.**
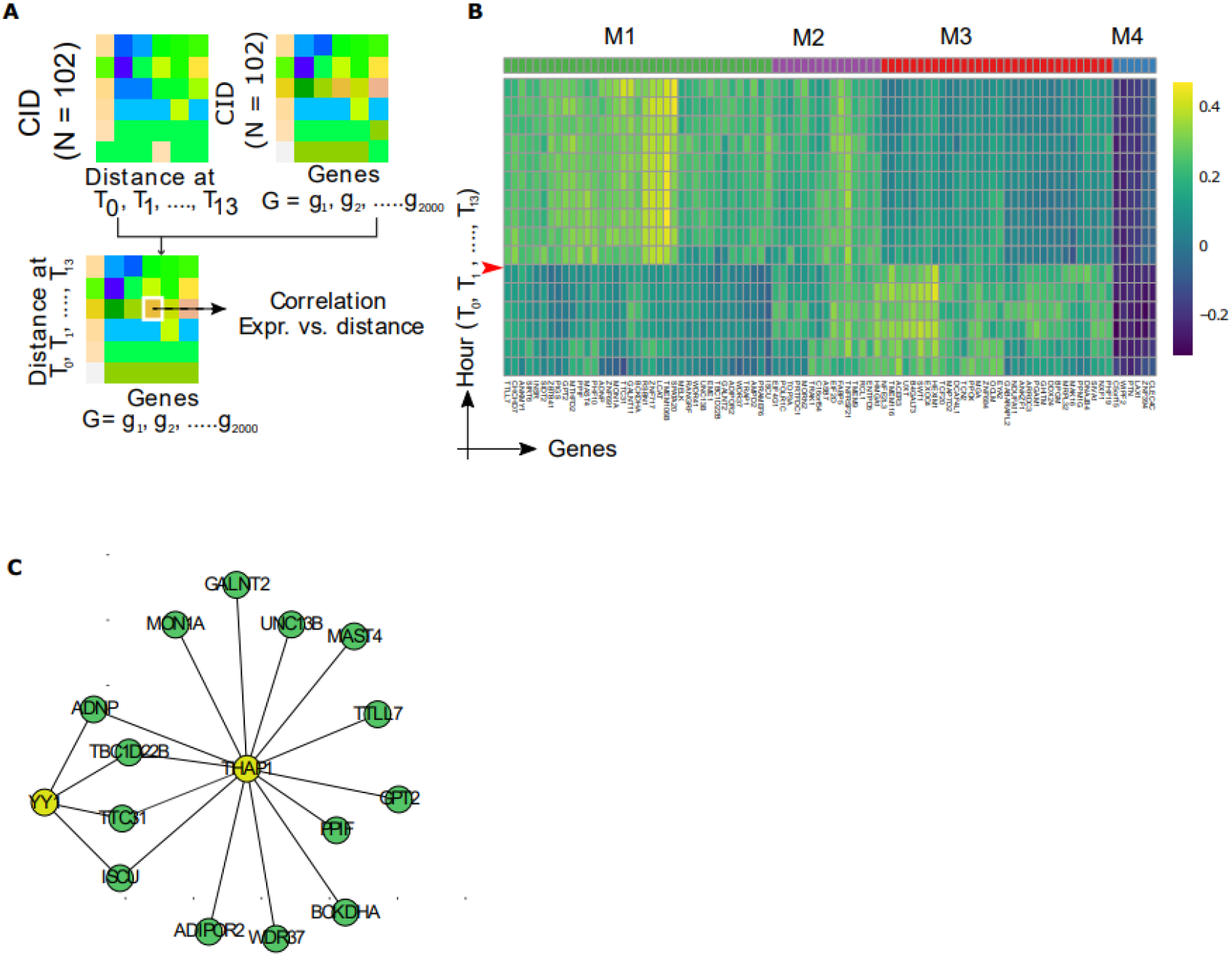
(A) Schematic workflow to estimate the correlation between cancer cell-NK cell (CIDs) distance and terminal expression profile of CIDs. This takes two matrices as input - the first one, containing cell-cell distance across different time points, and the second one contains the gene expression measurements across the CIDs. (B) Heatmap shows correlation between cell-cell distances and terminal CID gene expression profiles. Genes showing strong association with the cell-cell distances are clustered into four groups based on the correlation patterns (M1 to M4). The heatmap reveals that physical distances between NK and cancer cells effect gene expression profiles that gets carried forward to define future cellular states. The time point of first cell culture medium exchange is shown in red arrow (C) Transcription Factor binding motif Enrichment Analysis of M1 specific genes and annotations of these motifs to TFs using RcisTarget identifies THAP1 and YY1 as potential transcriptional regulators.

### Cancer-Immune Cell Killing Events

The microfluidic system allowed us to design experiments that tracked NK cell killing events and the associated transcriptomic signatures. We observed ten NK cell killing events across a total of 132 CID interactions. Differential expression analysis between CIDs subgroups showed unique gene expression signatures between NK killing and non-killing events. Among the 187 up-regulated genes in the NK killing group were *CASP8*, *SH3BP2*, *IGF-1*, *CNPY1* and *LMO1* (Fig. 1D). We applied PROGgene V2, a tool for prognostic implications measurement of genes^36^, to the Cancer Genome Atlas (Breast Invasive Carcinoma; TCGA-BRCA patients), and analyzed the impact of these 187 up-regulated genes on overall survival. 164 out of 187 up-regulated genes overlapped with TCGA-BRCA dataset from PROGgene V2. We observed a subtle survival advantage in the patients having a higher mean expression of this combined gene signature (Hazard ratio (HR)= 0.11, p<0.05) (Fig. 3)

**Fig 3.**
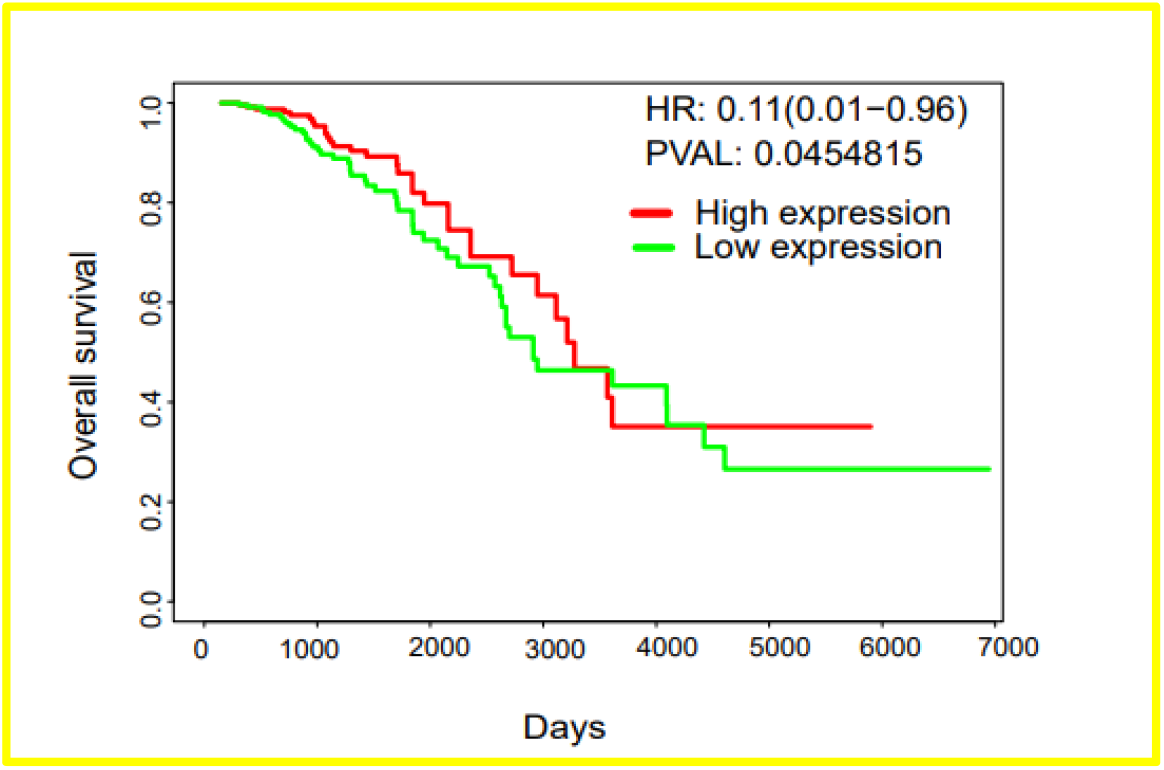
Kaplan-Meier overall survival plot for TCGA breast cancer (BRCA) cohort generated using PROGgeneV2. A gene signature associated with NK-killing cells event was averaged. To get gene signature value for each case, patients were classified into the low and high expression groups of the gene signature based on a median value with a P-value of 0.045 (Logrank test).

## Discussion

TNBC is a highly aggressive form of breast cancer with limited targeted treatment options. However, immunotherapy has recently turned out to be a promising strategy in the clinical management of the disease. In addition to current widely used T cell-mediated immunotherapy, research is also focused on NK cells that perform key roles in innate immune response in cancer. As such, comprehensive characterization of interacting NK and cancer cells at single-cell resolution might unravel actionable biomarkers and pathways involved in tumor growth and progression.

Here, we performed gene expression analysis of NK-cancer cell doublets resulting from the interacting distance between doublets measured over time. Statistical and bioinformatic investigation of the data enabled identification of gene expression signatures associated with successful and unsuccessful cancer cell lysis (killing events). Using TCGA gene expression profile data, we further confirmed that the identified signatures are linked to patient survival. Using single cell analysis, we also noted that the heterogeneity of the two cancer subclones confounded the process of dimension reduction by overshadowing the distinction of the NK cells’ identity.

### Cell interaction distances and transcriptional memory

We noted activation of exclusive, exposure time-bound, gene regulatory modules governing cell-cell physical proximity during the life cycle of NK-cancer cell interaction under co-incubation. We also noted an association between CID cell distance and their terminally profiled transcriptome in live cells. Similarly, Gide and colleagues reported a potential association between cancer/immune cell proximity with anti-PD-1 therapy response in melanoma patients^37^. This underscores the importance of cell-cell distance as an informative parameter to understand immunosurveillance and response in cancer. We observed changes in gene expression modules over time that correlated to the distance between live cells, suggesting the existence of transcriptional memory. Recent work has highlighted the controlled synthesis and degradation of mRNA transcripts as a major regulatory strategy influencing cell fate decisions^38^.

Transcriptional memory is a phenomenon that allows cells to retain reversible memory to respond to similar stimuli encountered in the future^39^. At time point 6 (after five-hour incubation and a change in culture medium), we noted a new set of genes expressed. During the initial hours of co-incubation, the cells might undergo the formation of lytic synapses as reported previously^1,40^, post which their interactions might take a drift.

In our study, we observed a regulatory role of three potential transcription factors, namely *BRCA1*, *YY1*, and *THAP1*, in transcriptional memory. *BRCA1* is a well-studied tumor suppressor gene and has known implications in breast and ovarian cancers. Among the remaining two, *YY1* promotes oncogenic activities in breast cancer^41^, whereas *THAP1* plays a key role in DNA repair and is also found to be overexpressed in breast cancers^42^. One of the genes from module 1 is *TRAP1*, whose overexpression is involved in promoting breast tumor growth. On the other hand, it also suppresses metastasis by regulating mitochondrial dynamics^43^. Another gene *MELK* from this module promotes TNBC proliferation. Targeting *MELK* can result in cell cycle arrest by reducing cyclin B1 and increasing p27 and p-JNK^44^. *EYA2* is also involved in promoting breast cancer proliferation. Its overexpression results in an increase of proliferative markers cyclin E, PCNA, and EGFR^45^.

Cell-cell signaling is a major component of cancer-immune cell interactions. We used gene expressions as a surrogate for ligand-protein activities. We focused on cancer-specific ligand-protein interactions featured in the iTALK ligand-protein pairs database^46^. To estimate the extent of ligand-protein coordination, we computed Pearson’s correlation coefficient for molecule pairs across CIDs and the other single cells (Fig. 4). We observed an elevated correlation between *ANXA1* and EGFR in CIDs. *ANXA1* is involved in mediating endocytosis of the EGFR receptor ANXA1/S100A11 complex^47–49^, and mediates cell-cell communication via exosomal EGFR^50^. We also observed a higher correlation between CD24/SIGLEC10 in the CIDs. Similarly, higher coordination between EGFR and *HSP90AA1* was observed in CIDs.

**Fig 4.**
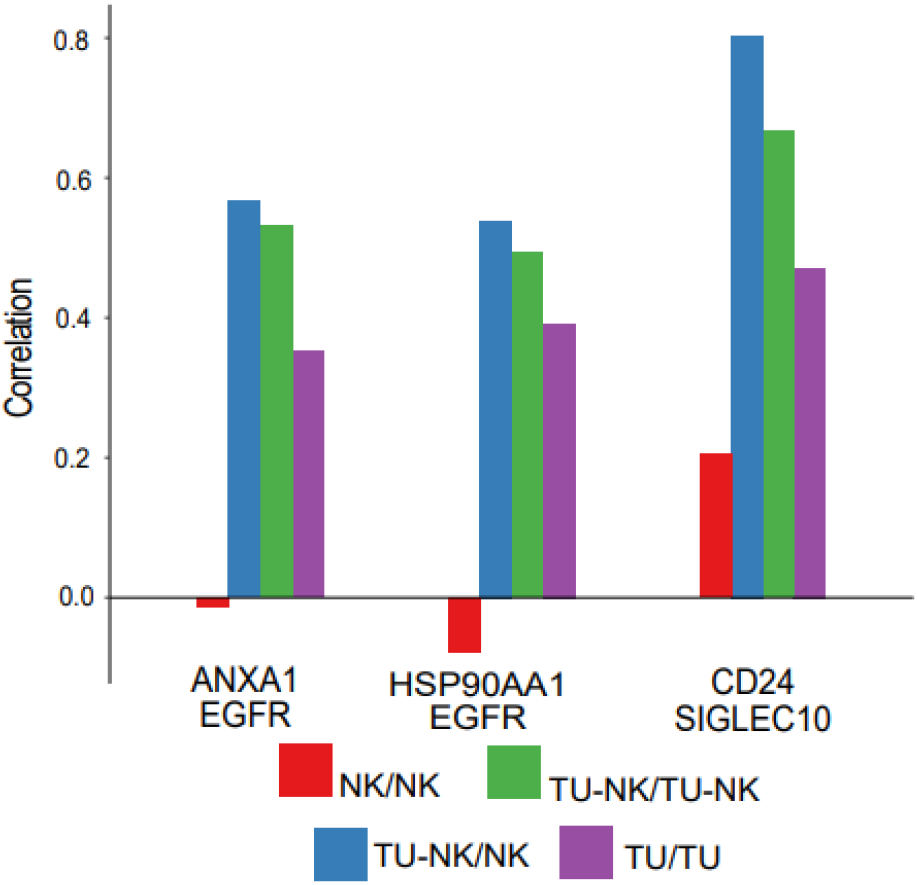
Impact of cell-cell signaling. Barplots showing an increased correlation in the CIDs for the three molecular pairs ANXA1/EGFR, HSP90AA1/EGFR, and CD24/SIGLEC10. NK = Natural Killer Cells; TU = Cancer/Tumor Cells. Legend key indicates cell status at the beginning and the end of the time-course tracking.

We observed preferentially elevated co-expression between CD24 (receptor) - SIGLEC10 (ligand) transcripts among NK-cancer doublets as compared to single NK and cancer cells. The coordination among this ligand-protein pair in CIDs suggests the importance of the CD24/SIGLEC10 axis in NK/TNBC cells. A past study reported the association of *SIGLEC10* in impeding NK cell function and poor patient survival in hepatocellular carcinoma (HCC); CD24/SIGLEC10 axis may thus be involved in regulating NK cell function^51^. In another study, the possibility of CD24/SIGLEC10 interaction in TNBC under the exposure of tumor-associated macrophages (TAMs) was reported. It has been observed that CD24 and SIGLEC10 are overexpressed in different tumor types and the TAMs, respectively^27^. Targeting this interaction may be therapeutically important. Another ligand-protein pair observed was the coordination between EGFR and *HSP90AA1*. *HSP90AA1* is critical for maintaining the stability and function of its client protein EGFR. This stabilization promotes pathogenesis in breast, head and neck cancer^52,53^. This occurs via EMT and tumor migration activating signaling pathways in MDA-MB-231 cells^54^.

### Cancer-Immune Cell Killing Events

When the CIDs were analyzed for cell killing events, we noted a gene expression signature that included *CASP8*, *SH3BP2, IGF-1,* and *LMO1. CASP8,* when activated through *FASLG,* results in activation of the extrinsic pathway of apoptosis in the target cells^55^. On the other hand, *SH3BP2^56^* and *IGF-1*^57^ have shown to play a decisive role in NK cell development and cytotoxicity. A strong differential expression signal was observed for several genes that are largely undocumented for their NK cytotoxicity links. These include the overexpression of *LMO1* (overexpressed in T lymphocytes from lymphoblastic leukemia)^58^, *CNPY1* and *ACSBG2* in the NK-killing group, which has not been described in NK cells till today.

Currently, it is difficult to co-incubate single cells in a controlled environment that simultaneously allows study of physical interactions between live cells and its effect on gene expression^59^. Here we report a cell-cell interaction study using an automated environment that precisely controls temperature, humidity, gas composition and media exchange to continuously monitor and measure the distance between cells. By processing the doublets on-IFC (in the same chambers) for cell lysis, reverse transcription, and cDNA amplification using a microfluidic multi-step chemistry, proximity measurements can be directly linked to downstream transcriptomics changes.

The use of this microfluidic system enabled the identification of unique features of NK cells’ anti-cancer activities. Highly coordinated gene expression profiles were identified as a result of dynamic changes in the physical distance of interacting NK and cancer cells, reinforcing transcriptional memory as a key regulatory strategy of cells. We could also trace increased coordination among some specific ligand-protein pairs, manifested through gene expression readouts. In the future, this microfluidic workflow could provide new leads when studying immuno-oncology cellular interactions. which may be considered while developing and administering NK cell-based immunotherapies.

## Data Availability

All raw and processed sequencing data generated in this study have been submitted to the NCBI Gene Expression Omnibus (GEO; https://www.ncbi.nlm.nih.gov/geo/) under accession number GSE181591 (https://www.ncbi.nlm.nih.gov/geo/query/acc.cgi?acc=GSE181591)

## Author contributions

Study Conception & Design: B.C.T.F., C.D.S., M.L., N.R., S.S.J.

Microfluidic System Development and Support: C.D.S., K.H., N.R.

Performed Experiment or Data Collection: B.C.T.F., C.D.S., N.M, P.K.K.

Computation & Statistical Analysis: S.C., N.M., N.R., D.S.

Data Interpretation & Biological Analysis: B.C.T.F., S.C., N.M, N.R., D.S., S.S.J.

Manuscript Writing, Review & Editing: B.C.T.F., S.C., N.M, L.T.D.C., N.R., D.S., S.S.J. with input from all the authors.

Supervision: L.T.D.C., N.R., D.S., S.S.J.

## Acknowledgement

N.M. and P.K.K. were supported in part by the John and Marva Warnock Research Fund.

S.S.J. is supported in part by the Stanford Catalyst for Collaborative Solutions.

B.C.T.F acknowledges the support of the Brazilian National Council for Scientific and Technological Development - CNPq; the Coordination for the Improvement of Higher Education Personnel (CAPES) (grant 88881.190291/2018-01) and the Coordination of A.C.Camargo - Antônio Prudente Foundation.

D.S. acknowledges the support of an intramural start up grant from the Indraprastha Institute of Information Technology, Delhi, and is partially supported by the INSPIRE faculty grant [DST/INSPIRE/04/2015/003068] from the Department of Science and Technology (DST), Govt. of India.

## Declaration of interests

K.H and N.R. are employees and stockholders of Fluidigm Corporation.

M.L is a former employee and stockholder of Fluidigm Corporation.

D.S. is a partner and equity holder at CareOnco Biotech Pvt. Ltd.

S.S.J. serves as a scientific advisor for Quantumcyte and Ravel Biotechnology.

## Material & Methods

### Standardization of NK cell activation

The activation of NK cells was established, in principle, by the evaluation of cell lines NK-92MI and MDA-MB-231. Activation was assessed after 24 and 48 hours of cell incubation in a proportion of 3 NK cells for each MDA-MB-231 by evaluating the markers (CD25, CD69, and CD314) expressed from cell-cell contact, using flow cytometry (Sony SH800S).

### Culturing MDA-MB-231 and NK-92MI

MDA-MB-231 cells (triple-negative breast carcinoma, ATCC® HTB-26 ™) were cultured using the DMEM (high modified) culture medium supplemented with 10% inactivated fetal bovine serum and 1% penicillin/streptomycin, replicating the culture every three days, keeping the culture at a confluence of 40% at the time of passage, at 37°C in a humid atmosphere at 5% CO2. The adherent cells were dissociated with the TrypLE reagent (Gibco) and resuspended in the complete culture medium.

NK-92MI cells (human NK cells genetically modified to produce interleukin 2, ATCC® CRL-2408) were cultured using the AMEM (Alpha Minimum Essential medium) culture medium, plus 0.2 mM inositol, 0.1 mM mercaptoethanol, 0.02 mM folic acid, 12.5% fetal bovine serum and 12.5% horse serum. The cells were homogenized to separate the clusters of NKs prior to the replication of the culture into a new 25 cm² flask (1mL of cells cultured previously + 9mL new culture medium). Both cell lines were cultured separately for the subsequent cell-cell interaction experiment using the Polaris system.

### Selection and incubation using the Polaris system (Fluidigm)

The first step is to prime the IFC with beads that will allow the adhesion of the cells to be incubated in the microchambers. After treatment, reagents and cells already labeled using specific markers to differentiate cell types (for NK cells, celltracker far red, and celltracker orange for cancer cells) and viability (calcein AM) were pipetted into the IFC. The selection of the cells to be incubated was performed on the Polaris system. It was configured for NKs and cancer cells to select positive cells for celltracker far red and calcein AM.

Some IFC wells were maintained with only one cell for later comparison of the gene expression of single cells with the gene expression obtained when the cell was incubated in doublets (NK cells + breast cancer cell lineage). After selection, the equipment was programmed to keep the cells in incubation for 16 hours, performing the replacement of culture medium (20% DMEM + 80% AMEM) every 5 hours, and time-lapse images every hour. Images were captured immediately before and after culture medium change.

71 single cancer cells, 77 single NK cells, and 132 cells in doublets (NK + cancer cells) were incubated, followed by the cell lysis, reverse transcription, and cDNA amplification. Subsequently, sequencing of the single-cell RNA, using NextSeq (Illumina) was performed.

### Distance measurement between cells

The distance shown in our study is the shortest distance between the membrane of MDA-MB-231 cell and NK cell on IFC within the doublets (NK + cancer cells) group. The distance has been measured for 13 time points. To analyze the videos frame by frame to yield the distance data, we used ImageJ, a public domain Java-based image processing software developed at the National Institutes of Health^60^.

### Data preprocessing

In total, we obtained expression profiles of 340 cells. The cells can be classified as follows. 1. Single NK cells (*n* = 77); 2. single tumor cells (*n* = 71); CIDs throughout all time points (*n* =132), and CIDs that were left with NK cells alone at the terminal time point (*n*=10). Out of the 340 cells, we removed 50 expression profiles. The cells discarded include — 1. 2 bulk RNA-Seq replicates of each of the NK-92MI and 2 MDA-MB-231 cell-lines; 4 empty chambers (no cells could be traced in the chambers from the start to the end); 8 empty chambers that initially contained tumor cells; 10 empty chambers that initially contained NK cells; 2 CIDs that started with single NK cells; 22 tumor cells that started as CIDs. After removal of these transcriptomes, we were left with 290 cells/doublets. Further, we discarded cells having less than 2000 expressed genes (non-zero RNA-seq by Expectation Maximization (RSEM) expression value^61^). Next, we retained genes having RSEM expression >5 in at least 10 cells. After these filtering steps, we were left with an expression matrix constituting 290 cells and protein-coding 8907 genes.

### Batch correction, clustering, and visualization of the single cells

The matrix obtained after performing the basic pre-processing steps was used as input for the Seurat, single cell analysis R package, as well as for other downstream analyses. We used Seurat’s data integration workflow to process the data with the genes detected in at least 5 cells, further employing the standard routines for log-normalization, variance stabilizing transformation for identification of highly variable genes using NormalizeData() and FindVariableFeatures() with default parameter settings. The cells used in this study originated from two independent runs. For integration and batch correction, we used the FinIntegrationAnchors() with k.filter=100 for identification of anchor cells that represent matching cell pairs across the two datasets in order to project the transcriptomes into a shared space and IntegrateData() function was used for integrating these anchors, which involve Canonical Correlation Analysis (CCA). These steps provided the batch corrected matrix. 2-D map of the cells was created using the RunUMAP() function.

### Differential gene expression analysis

Limma-voom^62^ was used to identify differentially expressed genes (DEGs) among the cell-groups. Top DEGs that qualified adjusted P-value cutoff of 0.05 and absolute log2 fold change cutoff of 1 were further analyzed for their biological significance.

### Survival analysis based on the upregulated genes governing NK cell antitumor activity

PROGgeneV2^36^ was used for gene signature based (based on a gene list) overall survival analysis combined gene signature analysis functionality using the TCGA-BRCA dataset, containing survival information for 594 patients. Out of 187 upregulated genes in NK cells showing antitumor activity, 164 genes were found overlapping with the TCGA-BRCA dataset. PROGgeneV2 produced a Kaplan-Meir plot, segregating the patients into high and low risk groups using the median value of the combined gene expression signature as cut-off.

### Regulation of cell-cell distance and transcriptional memory

We considered two matrices to track the association between temporally recorded cell-cell distance and terminally profiled gene expression. First, the processed gene expression matrix of dimensions |G| ×|C|, where |G|, and |C| denote the number of genes and the number of CIDs, respectively. Second, the cell-cell distance matrix of dimensions |C| ×|T|, where |T|denotes the time points when the cell-cell distance measurements were recorded. We used these two matrices to compute the third matrix of dimensions |G| ×|T| containing the Pearson’s correlation coefficients 〖ρ〗_ g,tfor each gene-time point pairs {(g,t) | t∈T,g∈G}. Notably, |G|=2000,|C|=102,and |T|=13. We retained 90 genes with 〖|ρ〗_ g,t|≥0.25for at least one time point for further analysis. These 90 genes were subjected to hierarchical clustering based on the computed correlation values. 4 gene modules were retrieved using cutree(). For each of these modules, motif enrichment and transcription factor activity analyses were performed using RcisTarget^35^ and ShinyGO^34^, respectively. RcisTarget discerned enriched TF binding motifs and reported a module-specific list of TFs, using the hg19-tss-centered-10kb-7species.mc9nr.feather database containing genome-wide ranking for the motifs. Gene regulatory networks were constructed using igraph^63^.

### Cell-cell signaling among CIDs

We used iTALK R package featuring information on 2,648 cancer-specific ligand-receptor interactions. We applied the rawParse() function to select the top 50% highly expressed genes from our processed gene expression matrix (constituting 290 cells and 8907 genes), using mean as a method of choice. This identified 230 ligand-receptor pairs using the FindLR() function. We computed Pearson’s correlation coefficient between expression values associated with the selected ligand-receptor pairs across CIDs (TU-NK/TU-NK, TU-NK/NK) and qualified ones that exhibited Pearson’s correlation coefficient > 0.4. To rule out the possibility that the co-expression stems solely from either NK or Tumor cells alone, we tracked Pearson’s correlation coefficient among NK/NK and TU/TU cases as well. With these criteria, 20 ligand-receptor pairs were selected, and three of these were found directly involved in breast cancer signaling.

## Supplementary Figures

**Fig S1.**
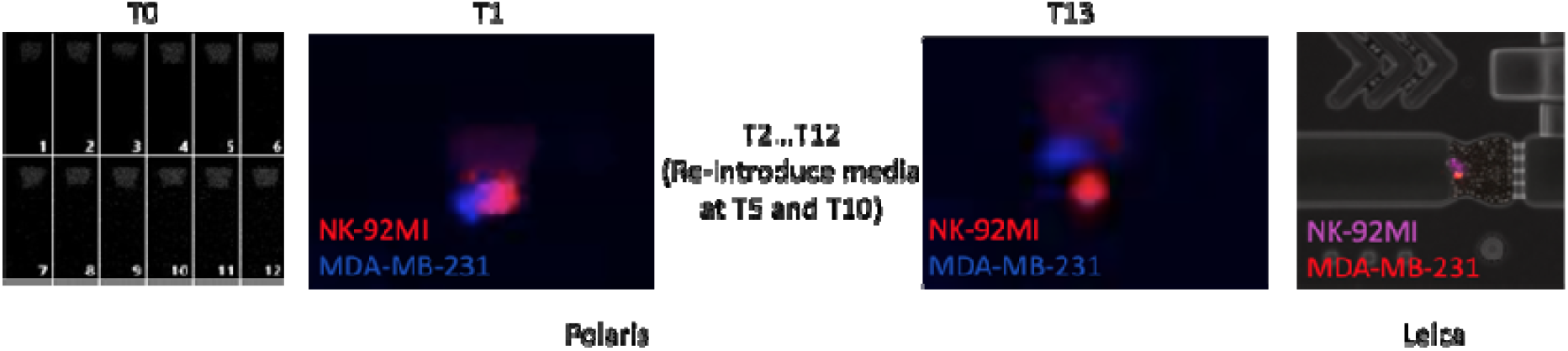
Single-cell or doublets culture and monitoring workflow. At the beginning of the experiment (T0), the NK cells and cancer cells were captured and imaged on the Polaris integrated fluidic circuit (IFC). The representative T0 figure shows cell culture chamber 1-12 out of 48 wells on one IFC. The cells were then incubated and monitored using Polaris system for 13 hours. Automated images were taken by Polaris system at one-hour interval (T1, T2, …T13). The NK-92MI cells are shown is pseudo red color, and MDA-MB-231 cells are shown in blue color. These time-lapse images were used for cell-cell distance measurement. The culture media were replenished every five hours automatically on the Polaris system. Before single cell RNA seq library preparation, high-resolution images were collected for each cell culture chamber using a Leica microscope system for viability test. The NK-92MI cells are color-coded in pink, and MDA-MB-231 cells are color-coded in red for the microscopic image.

**Fig S2.**
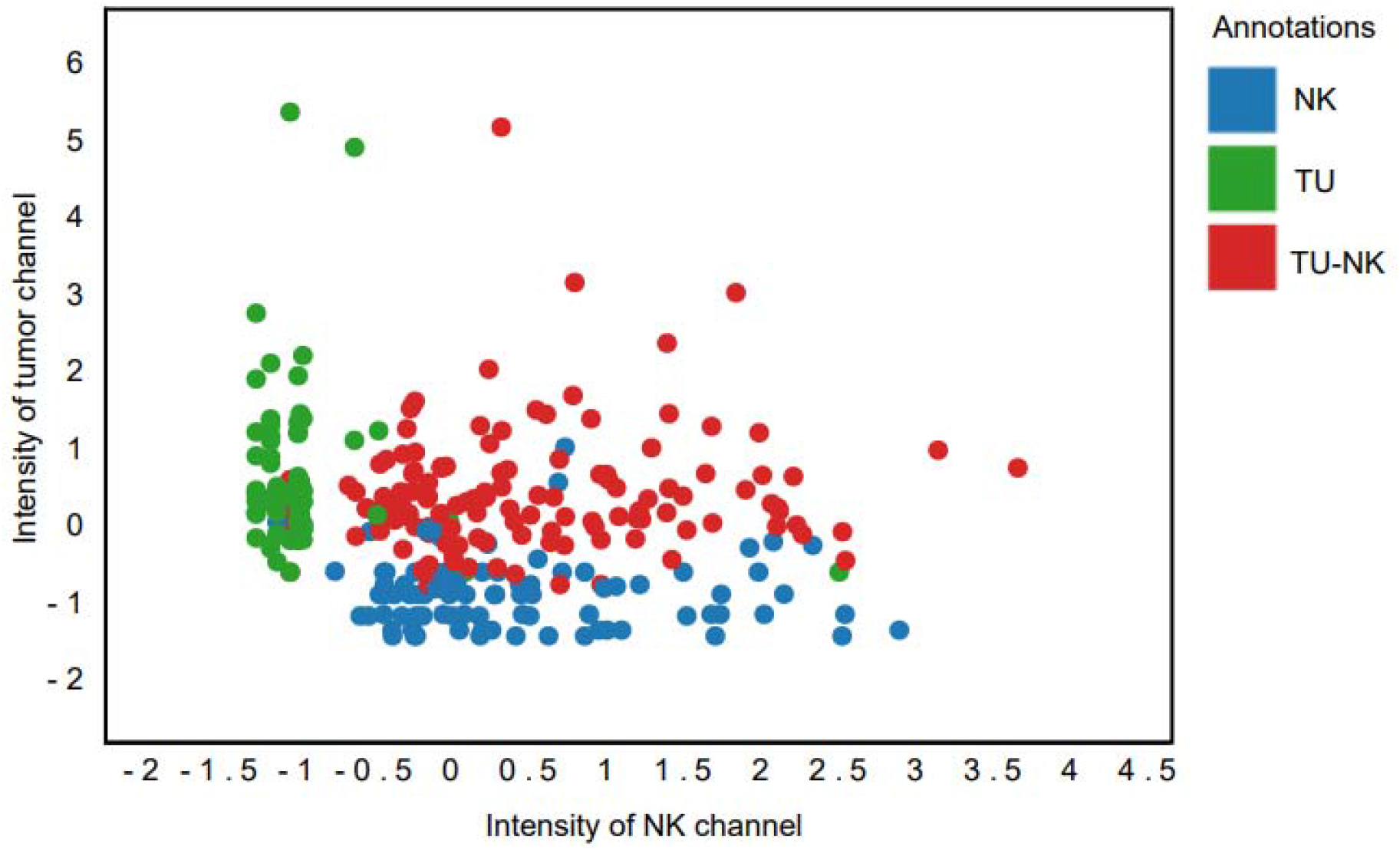
Scatterplot shows z-score normalized fluorescence intensities of NK and tumor imaging channels, and as per the grouping of cells, these corroborate with the original cell type annotations.

**Fig S3.**
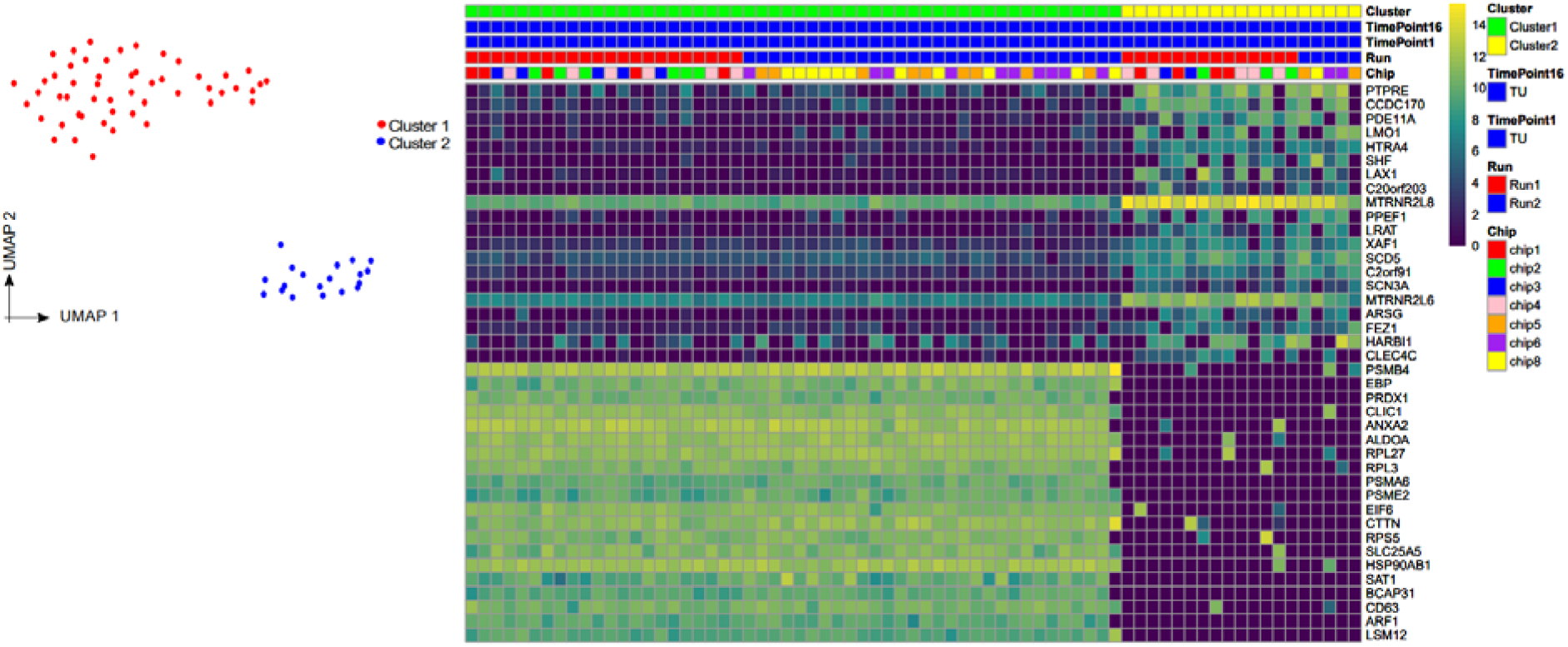
(A) UMAP based visualization of single tumor cells showing heterogeneous populations of tumor cell lines. (B) Heatmap showing differential genes between two tumor cell line clusters.

